# EIF3H Regulates ERK-Driven Oncogenic Signaling in Breast Cancer Metastasis

**DOI:** 10.1101/2025.08.24.672041

**Authors:** Shruti Bandyopadhyaya, Janvi Patel, Prashanthi Karyala, Himanshu Agrawal, Ekta Tripathi

## Abstract

Breast cancer remains a leading cause of cancer-related mortality among women, with metastasis being the primary driver of poor prognosis. The ubiquitin–proteasome system (UPS) is a central regulator of protein homeostasis, and its dysregulation is associated with multiple cancers. Within this system, deubiquitinating enzymes (DUBs), which remove ubiquitin moieties from target proteins and thereby modulate their stability and function, have emerged as attractive therapeutic targets. Eukaryotic initiation factor 3 subunit H (EIF3H), a JAMM family DUB, is overexpressed in multiple cancers and implicated in stabilizing oncogenic proteins. Using clinical transcriptomic datasets, we identified EIF3H as significantly upregulated in breast invasive carcinoma, with high expression correlating with poor patient outcomes. Functional assays demonstrated that EIF3H overexpression enhances proliferation, migration, and invasion of breast cancer cells, whereas its knockdown suppresses these traits. Mechanistically, EIF3H physically interacts with and deubiquitinates phosphorylated ERK (pERK), preventing its degradation and sustaining MAPK pathway activation. This represents the first report of pERK as a direct EIF3H substrate, revealing a novel mechanism linking EIF3H to metastatic progression. Moreover, EIF3H-deficient cells display increased sensitivity to chemotherapeutic drugs, suggesting that pharmacological inhibition of EIF3H may simultaneously impair metastasis and improve therapeutic efficacy. Collectively, our findings identify EIF3H as a potential therapeutic target for combating metastatic breast cancer.

## Introduction

Breast cancer remains one of the most prevalent cancers worldwide and a leading cause of cancer-related mortality among women, with metastasis being the primary contributor to its high mortality rate [1]. The ubiquitin-proteasome system (UPS) plays a crucial role in protein homeostasis and is implicated in various cancers, including breast cancer [2]. The UPS regulates protein turnover, stability, and degradation by tagging target proteins with ubiquitin molecules, marking them for destruction by the 26S proteasome. Through this mechanism, governs key cellular processes such as the cell cycle, apoptosis, and the DNA damage response, making it an attractive target for cancer therapy [3]. Deubiquitinating enzymes (DUBs) counterbalance the UPS by removing ubiquitin from substrates, thereby influencing tumor initiation and progression [4]. Emerging evidence indicates that specific DUBs are dysregulated in cancer and contribute to cancer specific traits such as proliferation, invasion, metastasis, and therapy resistance [5, 6]. Thus, targeting DUBs offers a promising therapeutic strategy to modulate UPS activity in cancer, with significant potential in breast cancer treatment [7, 8].

Eukaryotic initiation factor 3 (eIF3) is the largest translation initiation complex, comprising 13 subunits and essential for multiple initiation steps including ternary complex recruitment, mRNA attachment, and scanning for the initiation codon. Among the subunits, EIF3H and EIF3F contain MPN domains that belong to the JAMM family of DUBs [9]. EIF3H has been found to be overexpressed in several cancers such as lung, prostate, colorectal, and breast cancer [10–12]. It was experimentally shown to possess DUB activity, notably by deubiquitinating and stabilizing YAP, thereby promoting tumor progression and metastasis in breast cancer [12]. EIF3H exerts oncogenic effects through the modulation of key cancer-related signaling pathways including EMT, Hippo, and Wnt/β-catenin [12–14] and stabilizing several proteins such as Myc, Snail, Hax1, and YAP through its DUB activity, driving tumor progression. Therefore, unraveling the molecular mechanisms underlying EIF3H function and mapping its downstream targets may provide valuable opportunities for the development of targeted therapies against metastatic breast cancer.

The mitogen-activated protein kinase (MAPK) pathway is a well-established driver of tumor progression and metastasis in a variety of cancers, including breast cancer [15, 16]. This pathway regulates essential cellular processes such as proliferation, survival, differentiation, and migration, all of which contribute to the metastatic cascade. The MAPK pathway is an evolutionarily conserved signaling cascade that regulates cellular responses through a three-tier kinase module, primarily controlled by phosphorylation and interactions with regulatory proteins. Emerging evidence highlights the role of UPS in modulating the stability and turnover of MAPK pathway components influencing signaling outcomes. In particular, ERK2, a member of the MAPK family, is ubiquitinated by the E3 ligase activity of MEKK1 and stabilized by deubiquitination through USP15 [17, 18]. Understanding UPS-mediated regulation of MAPK signaling may provide new avenues to better understand pathway dynamics and develop alternate methods to overcome resistance to kinase inhibitors.

Our previous work investigated the role of DUBs in breast cancer metastasis using publicly available clinical transcriptomic datasets and identified EIF3H as a candidate DUB of interest [19]. EIF3H was upregulated in breast invasive carcinoma, with high EIF3H expression strongly correlated with poor clinical outcomes. This upregulation was further validated in breast cancer cell lines and protein-protein interaction analysis revealed that EIF3H associates with several metastasis-related genes, suggesting its potential role in promoting breast cancer progression.

Building on these findings, the present study investigates the functional role of EIF3H in breast cancer metastasis. We demonstrate that EIF3H overexpression significantly enhances proliferation, migration, and invasion, while EIF3H knockdown produces the opposite effect. Mechanistically, EIF3H physically interacts with and deubiquitinates phosphorylated ERK (pERK), thereby stabilizing it and sustaining MAPK pathway activation, which in turn promotes metastatic behaviour. To our knowledge, this is the first report identifying pERK as a direct substrate of EIF3H, uncovering a previously unrecognized mechanism by which EIF3H contributes to breast cancer progression. Importantly, EIF3H-deficient cells exhibited increased sensitivity to standard chemotherapeutic agents, suggesting that small-molecule EIF3H inhibitors could be a promising strategy to block metastasis and enhance chemotherapeutic efficacy. Taken together, our findings establish EIF3H as a novel oncogenic regulator of MAPK signaling and a promising therapeutic target for suppressing breast cancer metastasis and improving chemotherapeutic responses.

## Materials and Methods

### Cell Lines and Culture Conditions

The T47D human breast cancer cell line was maintained in RPMI-1640 medium supplemented with 10% fetal bovine serum (FBS) and 1% penicillin–streptomycin (Pen/Strep) at 37 °C in a humidified 5% CO₂ atmosphere.

### Generation of EIF3H-Overexpressing and -Knockdown T47D Lines

T47D cells were transfected with N-Flag–tagged EIF3H expression plasmid (Sino Biologicals; HG16405-NF) using Lipofectamine 3000 Transfection reagent (ThermoFisher; #L3000015) as per the manufacturer’s instructions. Seventy-two hours post-transfection, cells were selected in medium containing 900 µg/mL hygromycin B (Gibco; #10687010) for three days. Resistant colonies were expanded and maintained in 400 µg/mL hygromycin B. Stable EIF3H overexpression was confirmed by qPCR and Western blot. For CRISPR-Cas9–mediated knockdown of EIF3H, two guide RNAs (gRNA1 and gRNA2) were designed and cloned into the lentiCRISPRv2 vector (https://www.addgene.org/crispr/zhang). The gRNA sequences are provided in Table 1. Lentiviral particles carrying EIF3H-targeting gRNA1, gRNA2, or a non-targeting control (NTC) were produced in HEK293T cells and used to transduce T47D breast cancer cells. Forty-eight hours later, cells were selected with 200 ng/mL puromycin (ThermoFisher; # A1113803) for 3 days. After selection, cells were maintained in 100 ng/mL puromycin. Knockdown efficiency was validated by qPCR and Western blot.

**Table 1:**
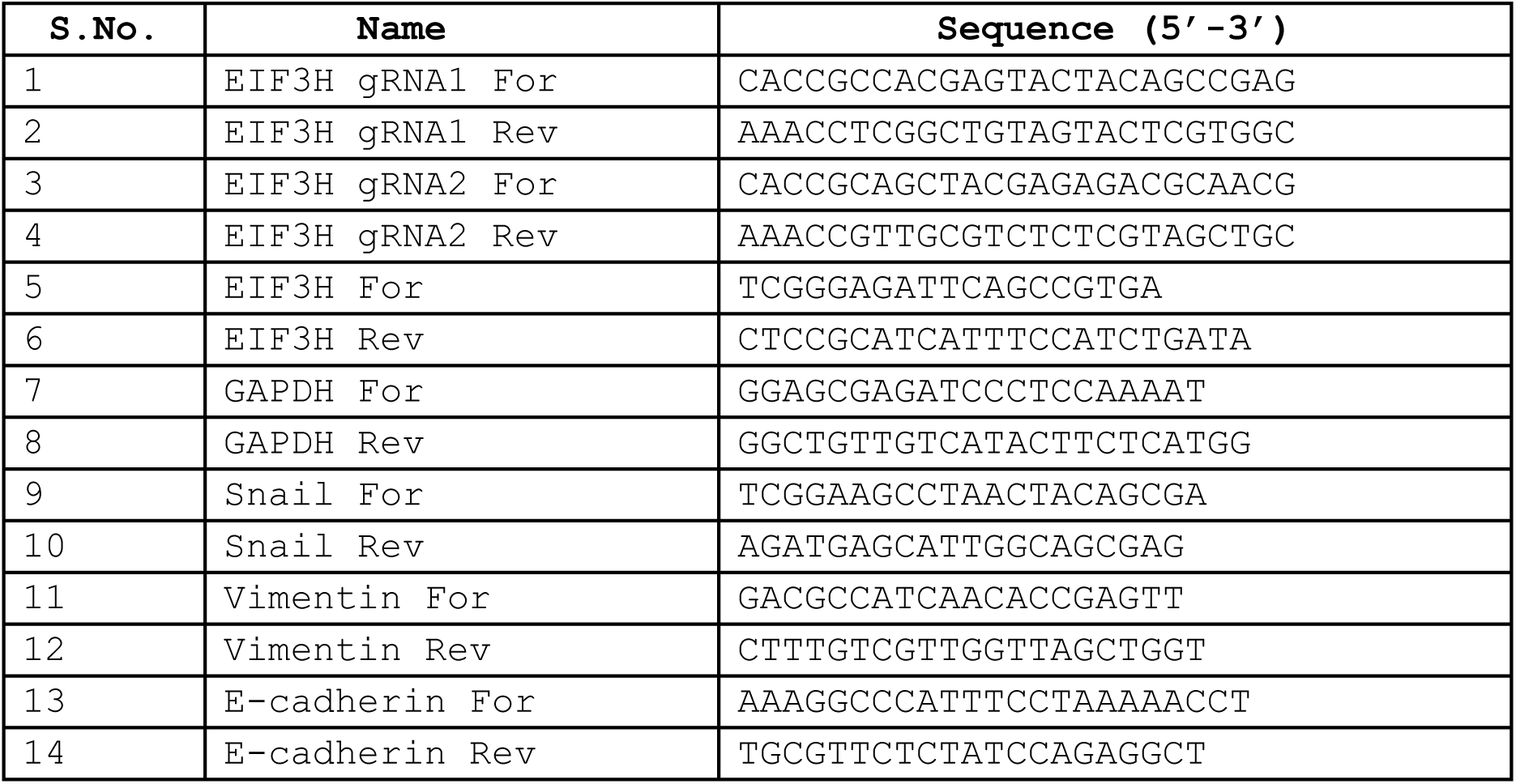
Primer Sequences.

### qPCR Analysis

Total RNA was isolated using the Qiagen RNeasy Mini Kit (# 74104). One microgram of RNA was reverse-transcribed into cDNA (Bio-Rad iScript™ cDNA Synthesis Kit; #1708891). Quantitative PCR was performed using 100 ng of cDNA per reaction using Bio-Rad iTaq™ Universal SYBR Green Supermix (#1725120), using primers specific for EIF3H and EMT markers (Snail, Vimentin, E cadherin), with GAPDH as the housekeeping gene. Reactions were run in duplicates. Relative gene expression was calculated by the ΔΔCt method, normalized to the housekeeping gene, and expressed as mean ± SEM from three independent experiments. The primer sequences are provided in Table 1.

### Cell Proliferation Assay

EIF3H-overexpressing and -knockdown T47D cells were seeded in 96-well plates, incubated at 37 °C in 5% CO₂, and proliferation was measured at 24 h, 48 h, 72 h, and 7 days. At each time point, CellTiter-Glo reagent (Promega #G7570) was added as per the manufacturer’s instructions. Briefly, plates were equilibrated for 20 min at room temperature, and luminescence was recorded on a microplate reader. All conditions were assayed in triplicate; raw luminescence values were normalized to the appropriate control sample, and proliferation curves were generated in GraphPad Prism (mean ± SEM).

### Clonogenic (Colony Formation) Assay

EIF3H-overexpressing and -knockdown lines, along with their controls, were trypsinized, counted, and plated in 12-well plates at densities optimized for each cell line. Triplicate wells per condition were maintained at 37 °C /5% CO₂ for 15-17 days, with medium changes every 3–4 days. At endpoint, wells were gently washed with PBS, fixed in methanol for 10 min, and stained with 0.5% crystal violet (Sigma # C0775) for 30 min. Excess dye was washed away with water, plates were air-dried, and colonies were counted using ImageJ. Colony counts were compared across groups to assess long-term proliferative capacity.

### Transwell Migration and Invasion Assays

For migration assays, 1×10⁶ EIF3H-overexpressing, knockdown, or corresponding control cells were suspended in 200 µL serum-free RPMI-1640 and added to the upper chamber using 8 µm PET Transwell inserts. For invasion assays, inserts were pre-coated with 200 µg/mL Matrigel (Sigma # 356234) and allowed to polymerize at 37 °C for 1 h. Following which, cells were then seeded in the coated inserts. In both assays, the lower chamber was filled with 600 µL RPMI-1640 containing 10% FBS as a chemoattractant. After 24 h, non-migrated or non-invaded cells were removed with a cotton swab and the remaining cells were labeled by incubating inserts in 4 µM Calcein-AM (Sigma #206700) diluted in non-enzymatic cell dissociation buffer (Sigma #C5914) for 30 min at 37 °C. The released fluorescence was measured on a microplate reader (excitation/emission 485/530 nm), and values were normalized to controls. Each condition was assayed in triplicate. For the ERK inhibitor experiments, EIF3H-overexpressing cells were treated with PD98059 (MedChem Express; #HY-12028,) at concentrations ranging from 5-60 µM for 1 hour, and transwell migration and invasion assay were performed subsequently.

### Wound-Healing (Scratch) Assay

EIF3H-overexpressing, knockdown, and control cells were seeded in 6-well plates to achieve ∼70-80% confluency. A uniform wound was created by scrapping the monolayer with a 200 µL pipette tip. The wells were gently rinsed with PBS to remove any debris and replenished with RPMI-1640 medium. Plates were incubated at 37 °C and 5% CO₂, and images of wound closure were captured at 0, 24, 48, and 72 h using a phase-contrast microscope (10X objective). Wound area was quantified with ImageJ software and expressed as the percentage of closure relative to the 0-h baseline. All conditions were performed in triplicate, and statistical significance was determined using Student’s t-test (GraphPad Prism).

### Western Blotting

EIF3H-overexpressing, knockdown, and control T47D cells were harvested and lysed in RIPA buffer supplemented with protease and phosphatase inhibitors and protein concentration was determined by BCA assay. 10 µg of total protein per sample was resolved by SDS–PAGE and transferred onto PVDF membranes (Millipore; #IPVH00010). Membranes were blocked in 5% BSA in TBST buffer for 1 h at room temperature, then incubated overnight at 4 °C with the following primary antibodies: [EIF3H (rabbit polyclonal; BioOrbyt orb539315; 1:1,000), phospho-ERK1/2 (rabbit mAb; CST 4370S; 1:1,000), total ERK1/2 (rabbit mAb; Abclonal A4782; 1:1,000), FLAG (mouse mAb; Sigma F3165; 1:1,000), GAPDH (rabbit mAb; CST 2118S; 1:5,000)] and HRP-conjugated secondary antibodies. The blots were washed thrice with TBST and incubated with the secondary antibody [anti-rabbit IgG (CST 7074S; 1:2,000) for all rabbit primaries, and anti-mouse IgG (CST 7076S; 1:2,000) for FLAG]. Blots were developed with Immobilon ECL Ultra substrate (Sigma #WBULS0500) and imaged on an ImageQuant system.

### Cycloheximide Chase Assay

To assess EIF3H protein stability, cells were treated with 50 µM cycloheximide (CHX) to inhibit de novo protein synthesis. At 0, 4, 16, and 24 h post-treatment, cells were lysed and 10 µg protein samples were processed for Western blotting as described above.

### Co-Immunoprecipitation (Co-IP)

To assess interactions between EIF3H and its target protein, EIF3H-overexpressing and vector control T47D cells were lysed in ice-cold TNE buffer (50 mM Tris-HCl pH 7.4, 150 mM NaCl, 1 mM EDTA) supplemented with protease inhibitors (Sigma #P8340). Lysates were precleared by incubation with Protein A/G agarose beads (Thermo # 20421) for 1 h at 4 °C. The supernatant was then incubated overnight at 4 °C with anti-FLAG antibody (Sigma #F3165) to capture FLAG-tagged EIF3H, followed by addition of fresh Protein A/G beads for 2 h. Beads were washed with cold TNE buffer to remove non-specific binders. Bound proteins were eluted by boiling in SDS–PAGE loading buffer, resolved on SDS-PAGE, and transferred to PVDF membranes. Blots were probed with FLAG,EIF3H, phospho-ERK, and GAPDH antibodies, then developed with ECL substrate. For the ubiquitination experiment, following Co-IP with the respective antibodies and SDS-PAGE, the blot was developed using an anti-ubiquitin antibody (CST # 3936).

### Drug Sensitivity Assay

To evaluate the effect of EIF3H knockdown on cell viability and proliferation, cells were seeded in 384-well plates. The following day, cells were treated with Doxorubicin (Sigma; # D1515) at concentrations ranging from 30 µM to 0.3 nM, and with Paclitaxel (Sigma; #T7402) at concentrations ranging from 30 nM to 0.3 pM. All treatments were maintained at a final concentration of 1% DMSO (Sigma; #2650) as a vehicle control. After 60 hours of incubation, cell viability was measured using the CellTiter-Glo Luminescent Cell Viability Assay (Promega #G7570). Luminescence was recorded using a plate reader, and data was analyzed using GraphPad Prism software to generate dose-response curves and calculate IC₅₀ values.

### Statistical analysis

Statistical analysis was performed using two-tailed unpaired or paired t-tests, and one-way ANOVA, as appropriate based on the number and type of groups being compared, using GraphPad Prism software. All experiments were independently conducted three times with replicates. Data are presented as mean ± SEM. A p-value of less than 0.05 was considered statistically significant.

## RESULTS

### 1. EIF3H Promotes Proliferation, Migration, and Invasion in Breast Cancer Cells

EIF3H plays a key role in the regulation of protein synthesis as part of the eukaryotic translation initiation complex and has been implicated in the development and progression of several cancers [5, 6]. Growing evidence, including studies from our group and others, highlights its role in breast cancer pathogenesis [12, 19]. To further investigate EIF3H function in breast cancer progression, we generated stable cell line models with both EIF3H overexpression and knockdown. T47D breast cancer cells were transfected with an EIF3H overexpression plasmid and selected with hygromycin. Surviving colonies were expanded, and EIF3H expression levels were measured using qPCR and Western blotting. qPCR analysis revealed a significant upregulation of EIF3H mRNA in the overexpression (OE) lines compared to vector controls (Fig. 1A). Consistently, Western blot analysis confirmed enhanced EIF3H protein levels, with Flag-tagged EIF3H detected in the OE cell lines (Fig. 1B & 1C), using GAPDH as a loading control.

**Figure 1:**
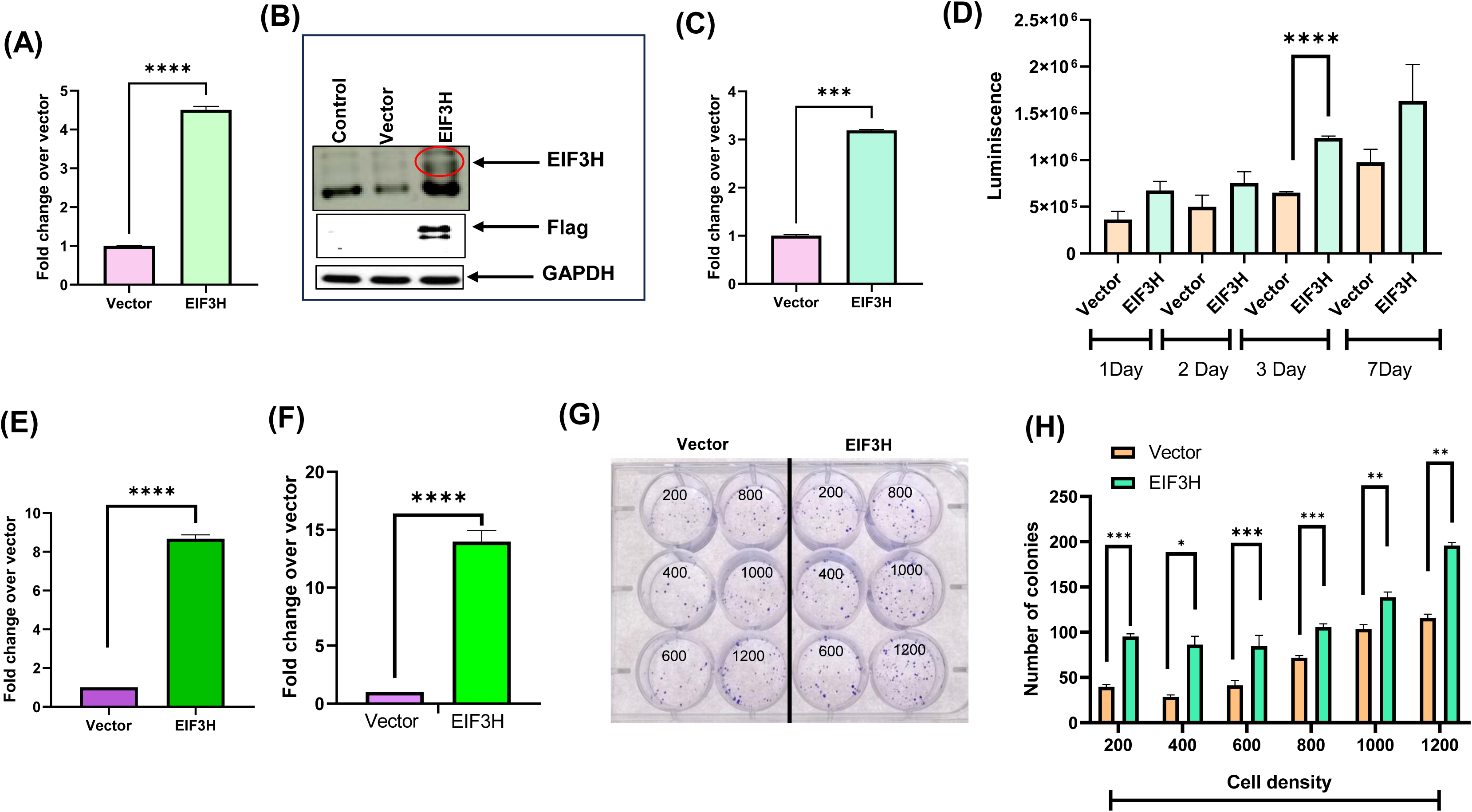
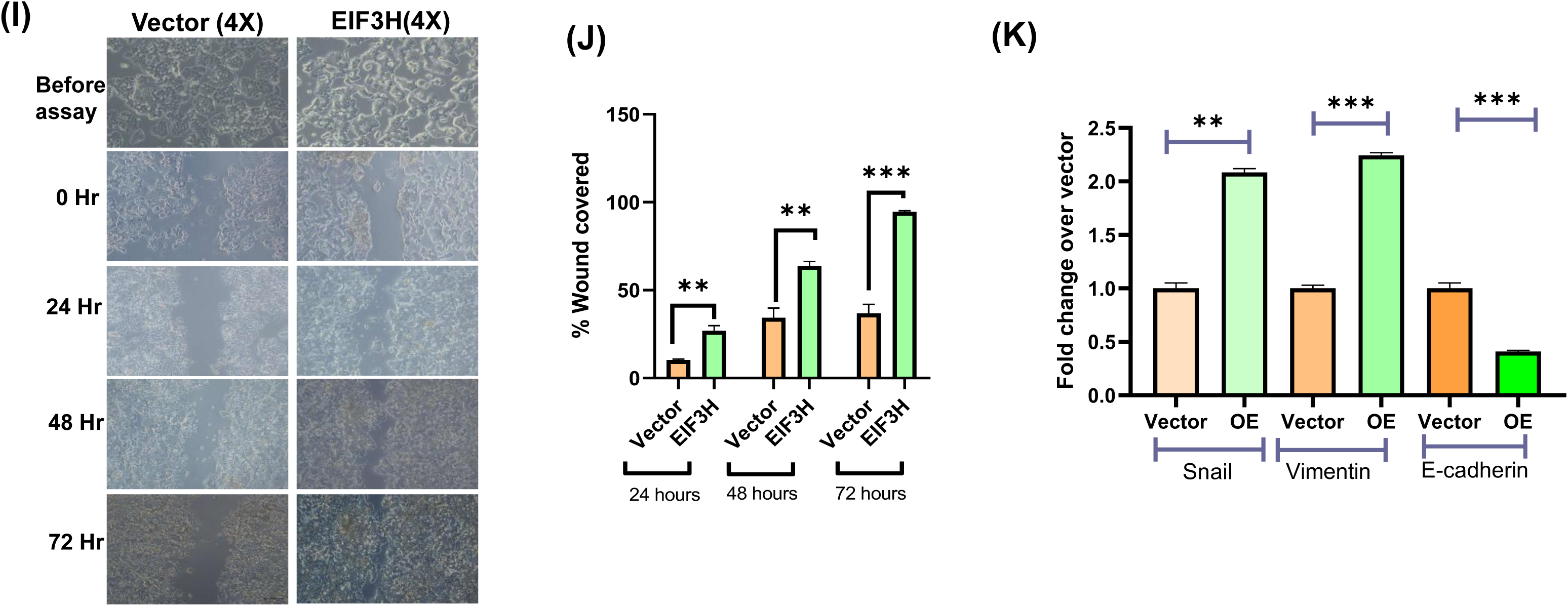
Functional Characterization of EIF3H Overexpression in Breast Cancer Cells. **(A)** Quantitative PCR: The mRNA expression levels of EIF3H were assessed in stable cell lines using EIF3H specific primers and normalised to a housekeeping gene GAPDH. **(B and C)** Western blot analysis: The protein expression of EIF3H was assessed using Flag antibody, with corresponding densitometric quantification. **(D)** Proliferation Assay: The proliferative capacity of EIF3H overexpression (OE) cells was assessed using the Cell Titer-Glo luminescence-based assay over a period of 7 days and luminescence was recorded at multiple time points to evaluate cell growth. **(E)** Transwell Migration Assay: Cell migration was examined using a Calcein AM-based **(F)** Invasion Assay: The invasive potential of EIF3H OE cells was evaluated using Matrigel-coated Transwell inserts, followed by Calcein AM staining **(G–H)** Clonogenic Assay: Long-term survival and clonogenic potential were evaluated by seeding EIF3H OE and vector control cells at increasing densities (200–1200 cells/well) and culturing them for 17 days. Colonies were stained with crystal violet and imaged to assess colony formation efficiency. **(I-J)** Wound Scratch Assay: To assess cell motility and wound closure capability, scratch assays were performed. The percentage of wound closure was monitored at 24, 48, and 72 hours in EIF3H OE and vector control cells. **(K)** qPCR : The mRNA expression levels of EMT markers (Snail, vimentin and E-cadherin) were assessed in stable cell lines using specific primers and normalised to a housekeeping gene. All data are presented as mean ± SEM from three independent experimental replicates. Statistical significance was determined as follows: *p < 0.05, **p < 0.01, ***p < 0.001, ****p < 0.0001.

To evaluate the role of EIF3H in breast cancer cell proliferation, we utilized EIF3H-OE cells and performed a CellTiter-Glo (CTG) viability assay over a period of 1, 2, 3, and 7 days. EIF3H OE cells displayed a increase in proliferation compared to vector control cells, with the significant difference observed on day 3 (Fig. 1D). Furthermore, transwell migration and invasion assays demonstrated that EIF3H OE cells exhibited increased migratory and invasive capabilities (Fig. 1E and F). To evaluate long-term proliferative capacity of EIF3H, we performed a clonogenic assays in EIF3H-OE cells. EIF3H-OE cells formed significantly more colonies compared to control cells, indicating an increased clonogenic capacity (Fig. 1G and H). A wound healing assay was also performed to assess cell motility and EIF3H OE cells showed accelerated wound closure, further supporting enhanced migratory behaviour (Fig. 1I and J). We next examined the expression of EMT (epithelial–mesenchymal transition) markers, including Snail, Vimentin, and E-cadherin using qPCR. As shown in Fig. 1K, Snail and Vimentin levels were upregulated, while E-cadherin expression was reduced, supporting the notion that EIF3H plays a role in promoting metastasis.

Conversely, the same set of assays was performed using EIF3H knockdown (KD) cells. To generate EIF3H-KD cells, two guide RNAs (gRNAs) targeting EIF3H and a non-targeting control (NTC) were cloned into the plentiCRISPRv2 vector and transduced into T47D cells. Stable knockdown lines were selected using puromycin. qPCR analysis confirmed a substantial reduction in EIF3H transcript levels, while Western blotting demonstrated significant knockdown of EIF3H protein expression (Fig. 2A-C). The same set of experiments were performed to assess the proliferation, migration and invasion of the EIF3H in knockdown status. EIF3H-KD cells displayed markedly reduced proliferation, migration, invasion, clonogenicity, wound healing activity and expression of Snail and vimentin compared to non-targeting controls (Fig. 2D-K). Collectively, these findings strongly support a pro-tumorigenic and pro-metastatic role for EIF3H in breast cancer cells.

**Figure 2:**
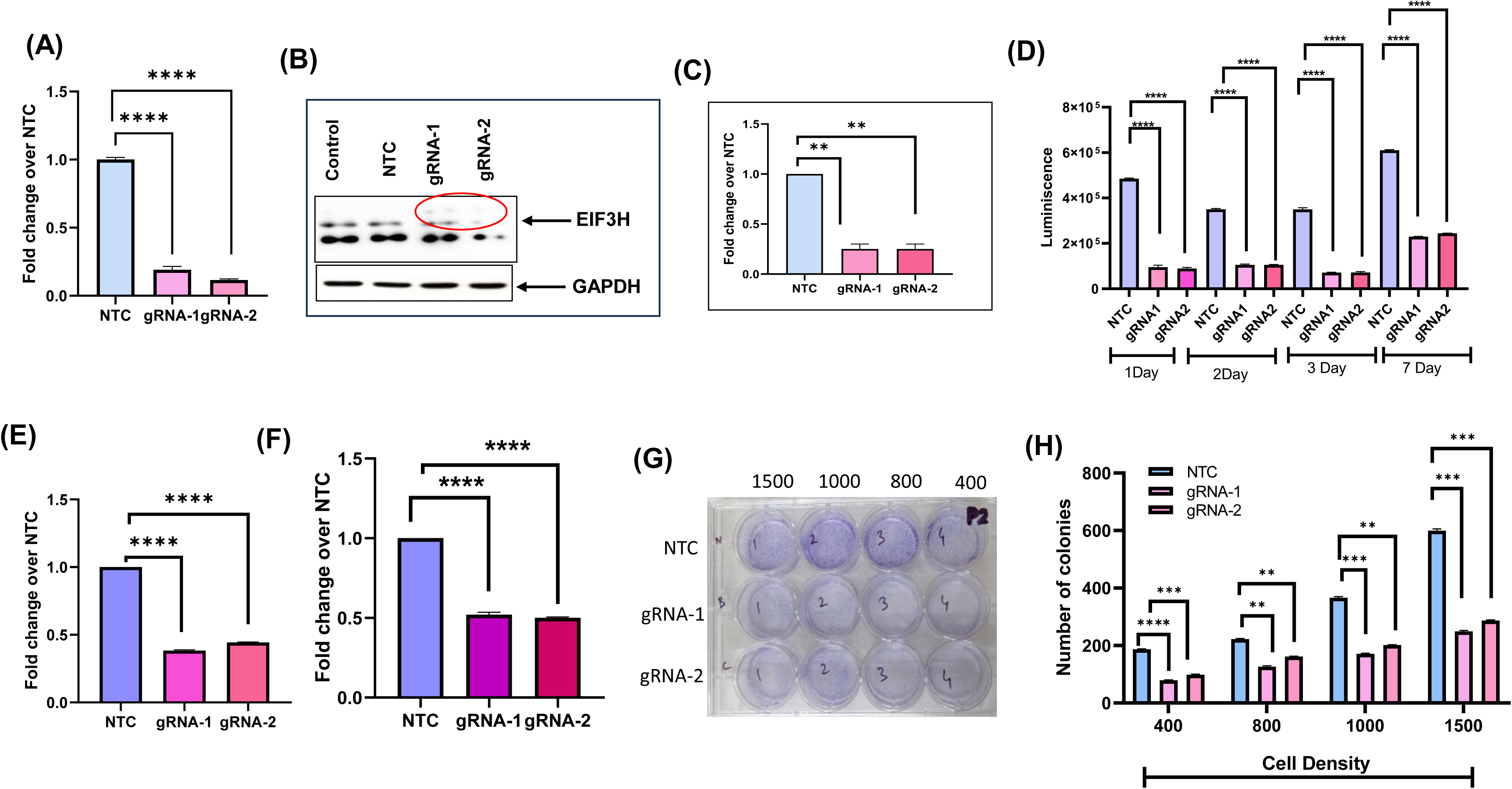
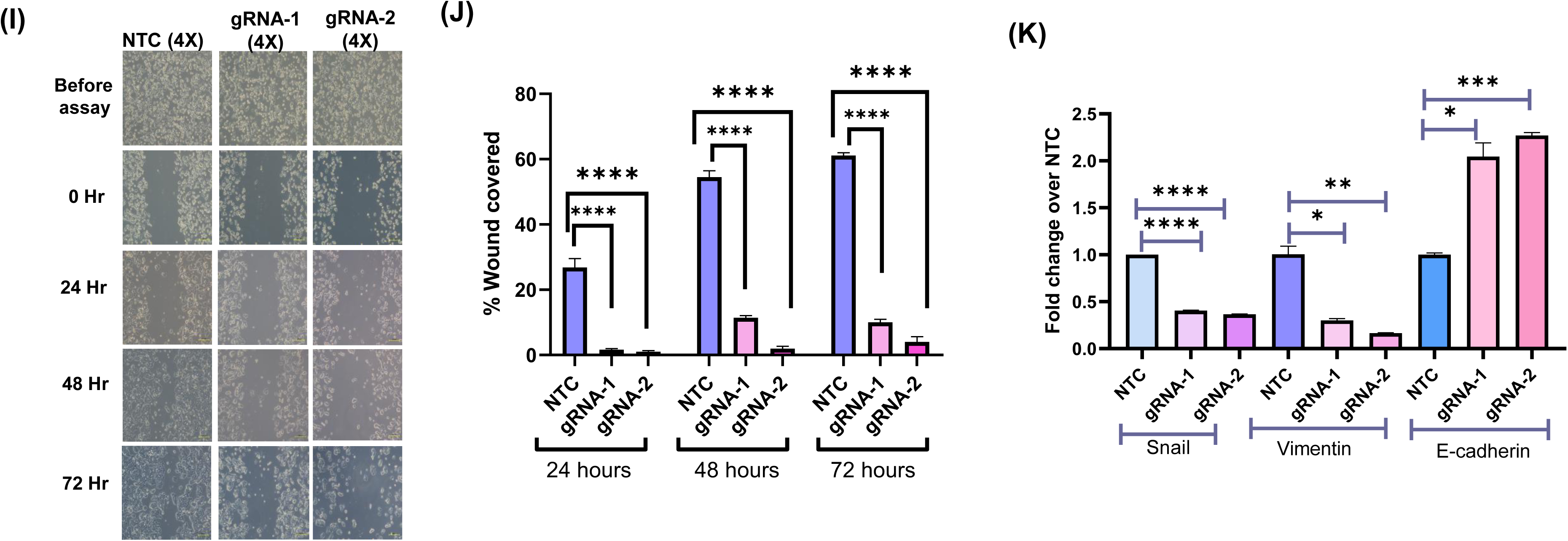
Functional Characterization of EIF3H Knockdown in Breast Cancer Cells. **(A)** Quantitative PCR: The mRNA expression levels of EIF3H were assessed in knockdown stable cell lines using EIF3H specific primers and normalised to a housekeeping gene. **(B and C)** Western blot analysis: The protein expression was assessed using EIF3H antibody, with corresponding densitometric quantification. **(D)** Proliferation Assay: The proliferative capacity of EIF3H Knockdown (KD) cells was assessed using the Cell Titer-Glo luminescence-based assay over a period of 7 days and luminescence was recorded at multiple time points to evaluate cell growth. **(E)** Transwell Migration Assay: Cell migration was examined in EIF3H NTC and KD cells using a Calcein AM-based Transwell migration assay **(F)** Invasion Assay: The invasive potential of EIF3H KD cells was evaluated using Matrigel-coated Transwell inserts, followed by Calcein AM staining **(G–H)** Clonogenic Assay: Long-term survival and clonogenic potential were evaluated by seeding EIF3H KD and NTC control cells at increasing densities (400–1500 cells/well) and culturing them for 17 days. Colonies were stained with crystal violet and imaged to assess colony formation efficiency. **(I-J)** Wound Healing Assay: To assess cell motility and wound closure capability, scratch assays were performed. The percentage of wound closure was monitored at 24, 48, and 72 hours in EIF3H KD and NTC cells. **(K)** qPCR : The mRNA expression levels of EMT markers (Snail, vimentin and E-cadherin) were assessed in stable cell lines using specific primers and normalised to a housekeeping gene. All data are presented as mean ± SEM from three independent experimental replicates Statistical significance was determined as follows: *p < 0.05, **p < 0.01, ***p < 0.001, ****p < 0.0001.

### 2. EIF3H Regulates MAPK Signaling by Stabilizing pERK

To explore the potential role of EIF3H in regulating oncogenic signaling pathways, we examined the activation status of MAPK pathway in T47D cells with altered EIF3H expression. Given the central role of the MAPK pathway in promoting cell proliferation, survival, and tumor progression, we investigated whether EIF3H regulates this signaling pathway. Western blot analysis revealed that EIF3H overexpression resulted in a marked increase in phosphorylated ERK (p-ERK) levels, indicating activation of the MAPK signaling cascade (Fig. 3A and 3B). In contrast, EIF3H knockdown significantly reduced p-ERK levels compared to control cells (Fig. 3C and 3D), suggesting that EIF3H can activate ERK phosphorylation. Notably, total ERK levels remained unchanged in both conditions, indicating that EIF3H specifically regulates ERK phosphorylation rather than its overall expression.

**Figure 3.**
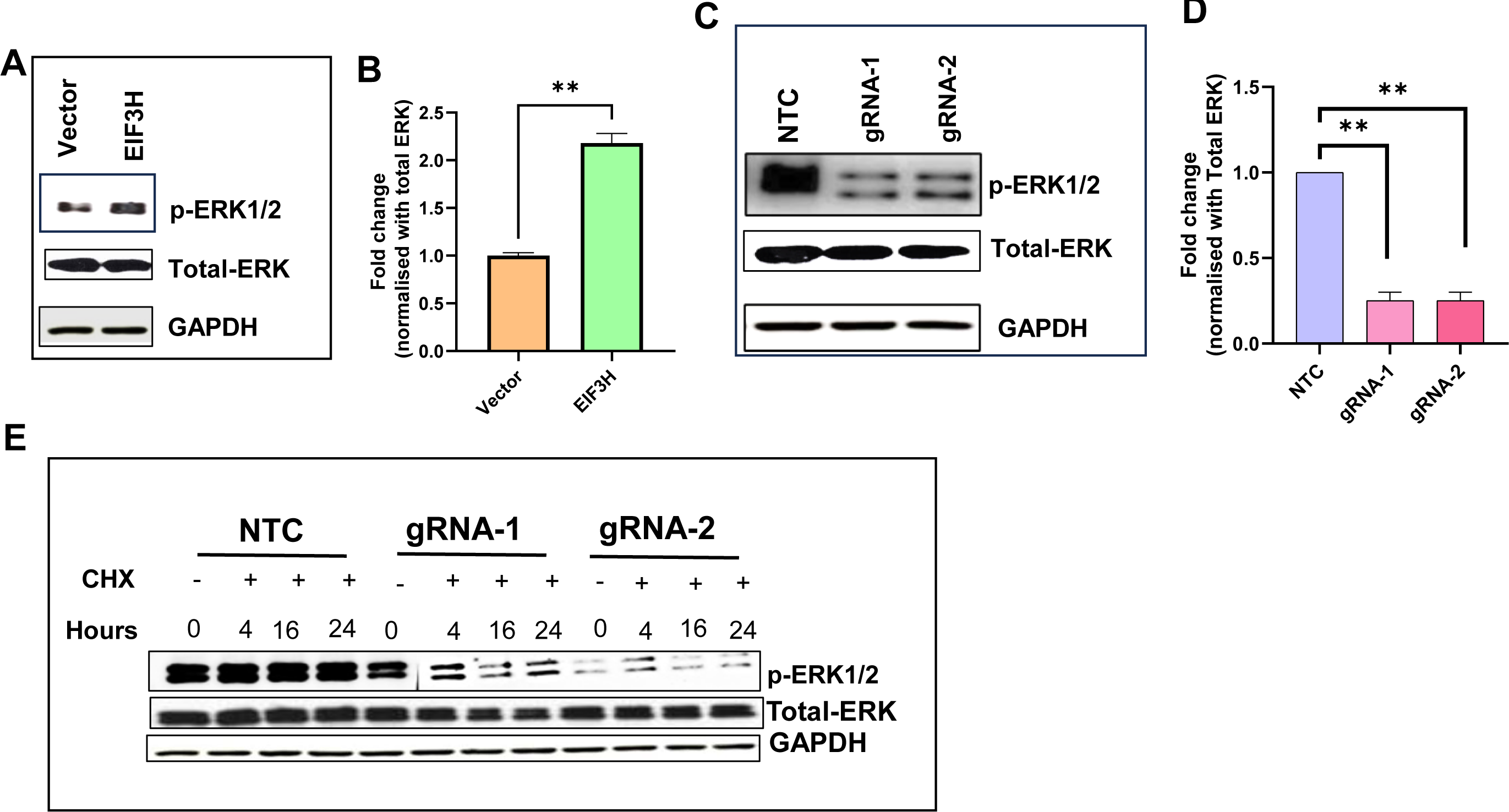
EIF3H Modulates p-ERK/MAPK Signaling Pathway and Stabilization: **(A–B)** Western blot analysis showing p-ERK levels in EIF3H-overexpressing cells. **(C–D)** Western blot analysis showing p-ERK1/2 levels in EIF3H knockdown cells. GAPDH and total ERK were used as loading controls.**(E)** Cycloheximide (CHX) chase assay: EIF3H knockdown cells were treated with CHX for the indicated time points (0–24 h), and p-ERK levels were assessed by western blot. Total ERK and GAPDH are included as loading controls.

To further confirm whether EIF3H specifically affects the stability of p-ERK, we performed a cycloheximide chase assay. Cycloheximide is a translational inhibitor that halts new protein synthesis, allowing us to monitor the degradation kinetics of existing proteins over time. EIF3H knockdown cells and control cells were treated with cycloheximide, and the levels of p-ERK were assessed at various time points post-treatment. Strikingly, EIF3H-deficient cells showed a marked, time-dependent decrease in p-ERK levels compared with controls, indicating accelerated degradation of p-ERK in the absence of EIF3H (Fig. 3E). In contrast, levels of total ERK remained largely unchanged, demonstrating that EIF3H selectively stabilizes the phosphorylated form rather than influencing overall ERK stability. Together, these findings establish EIF3H as a critical regulator of MAPK/ERK signaling by deubiquitinating and protecting p-ERK from proteasomal degradation. This selective stabilization of active ERK highlights a previously unrecognized mechanism by which EIF3H sustains oncogenic signaling, underscoring its potential as a therapeutic target in ERK-driven cancers.

### 3. EIF3H Interacts with and Regulates the Ubiquitination of ERK

Since EIF3H overexpression led to enhanced p-ERK signaling, we hypothesized that EIF3H may directly interact with p-ERK. To test this, we performed co-immunoprecipitation (Co-IP) assays. Lysates from EIF3H-overexpressing and vector control cells were immunoprecipitated with an anti-FLAG antibody to pull down EIF3H and its associated proteins. Subsequent Western blotting with anti-pERK antibodies revealed a clear interaction between EIF3H and p-ERK in the OE cells, whereas no interaction was observed in vector controls (Fig. 4A). Interestingly, EIF3H also associated with total ERK, suggesting that EIF3H may bind ERK irrespective of its phosphorylation status. These findings demonstrate a physical association between EIF3H and p-ERK, providing a mechanistic basis for how EIF3H sustains MAPK signaling and promotes various oncogenic processes.

**Figure 4.**
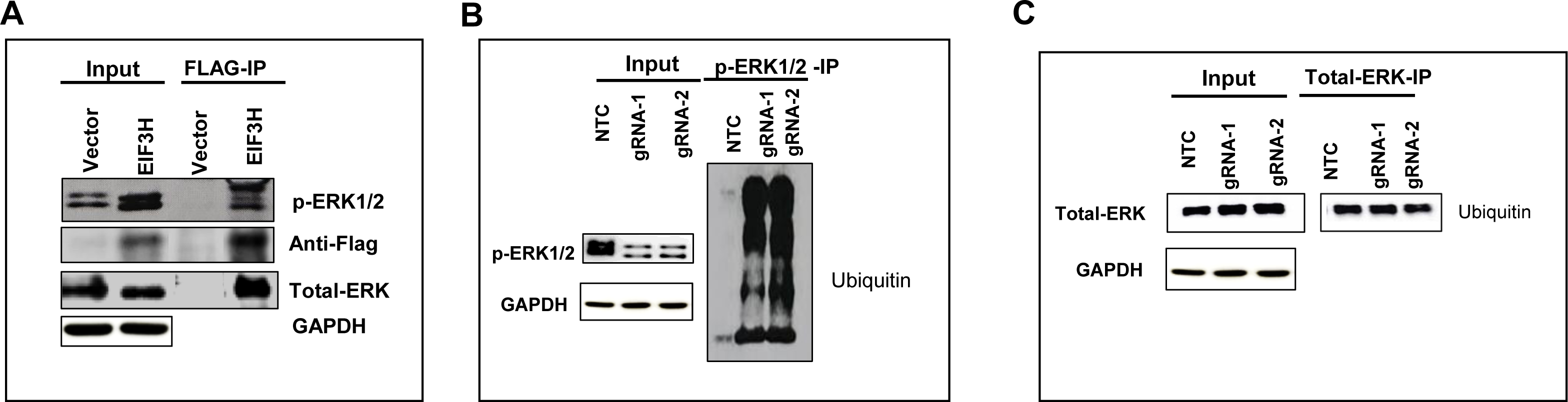
EIF3H Interacts with and Regulates Ubiquitination of pERK. **(A)** Co-immunoprecipitation (Co-IP) Assay: Lysates from EIF3H-overexpressing cells were immunoprecipitated using an anti-FLAG antibody. The precipitated complexes were analyzed by Western blot using anti-Flag, anti-pERK1/2 and total ERK antibodies. Input lysates were also probed with the same antibodies to assess expression levels. **(B)** Ubiquitination Assay: Co-IP was performed in EIF3H KD and NTC cells using an (B) anti-pERK1/2 antibody and **(C)** anti-Total ERK antibody, followed by Western blotting with an anti-ubiquitin antibody to assess ERK ubiquitination.

To further investigate whether EIF3H deubiquitinates pERK, we performed Co-IP assays in EIF3H KD cells. Immunoprecipitation using an anti-pERK antibody followed by Western blotting for ubiquitin revealed prominent ubiquitin smears in EIF3H KD cells, indicating increased ubiquitination of pERK in the absence of EIF3H (Fig. 4B). In contrast, NTC cells showed minimal or no pERK ubiquitination. These results suggest that EIF3H regulates pERK stability by preventing its ubiquitination. In EIF3H-deficient cells, enhanced ubiquitination likely targets pERK for proteasomal degradation, contributing to reduced pERK levels and impaired downstream signaling. Notably, when we performed immunoprecipitation using a total ERK antibody and probed for ubiquitin, no significant ubiquitination smear was observed (Fig. 4C). This finding indicates that EIF3H specifically deubiquitinates and stabilizes pERK, rather than total ERK. The reduction in pERK stability may underlie the observed decrease in proliferation, migration, and invasion in EIF3H KD cells.

To determine whether EIF3H directly regulates pERK ubiquitination, we performed Co-IP assays in EIF3H KD cells. Immunoprecipitation with an anti-pERK antibody followed by Western blotting for ubiquitin revealed prominent ubiquitin smears in EIF3H KD cells, indicating increased pERK ubiquitination in the absence of EIF3H (Fig. 4B). By contrast, NTC cells displayed minimal ubiquitination, suggesting that EIF3H protects pERK from ubiquitin-mediated degradation. Consistent with this, immunoprecipitation with a total ERK antibody followed by ubiquitin blotting showed no significant ubiquitination signal (Fig. 4C), confirming that EIF3H specifically targets the phosphorylated form. These findings demonstrate that while EIF3H can associate with both total ERK and pERK, it specifically deubiquitinates and stabilizes the phosphorylated form. By protecting pERK from ubiquitin-mediated proteasomal degradation, EIF3H sustains MAPK pathway activity. Conversely, loss of EIF3H leads to increased pERK ubiquitination, accelerated degradation, and impaired downstream signaling, which may explain the reduced proliferation, migration, and invasion observed in EIF3H-deficient cells.

### 4. ERK Signaling Modulates EIF3H-Mediated Cell Migration, Invasion, and Proliferation

To further probe the functional relationship between EIF3H and pERK, we treated EIF3H-overexpressing cells with PD98059, a selective MEK inhibitor that blocks ERK phosphorylation. Inhibition of ERK activity markedly reduced both migration and invasion (Fig. 5A and B), suggesting that the pro-migratory and pro-invasive effects of EIF3H are at least partially dependent on ERK pathway activation.

**Figure 5.**
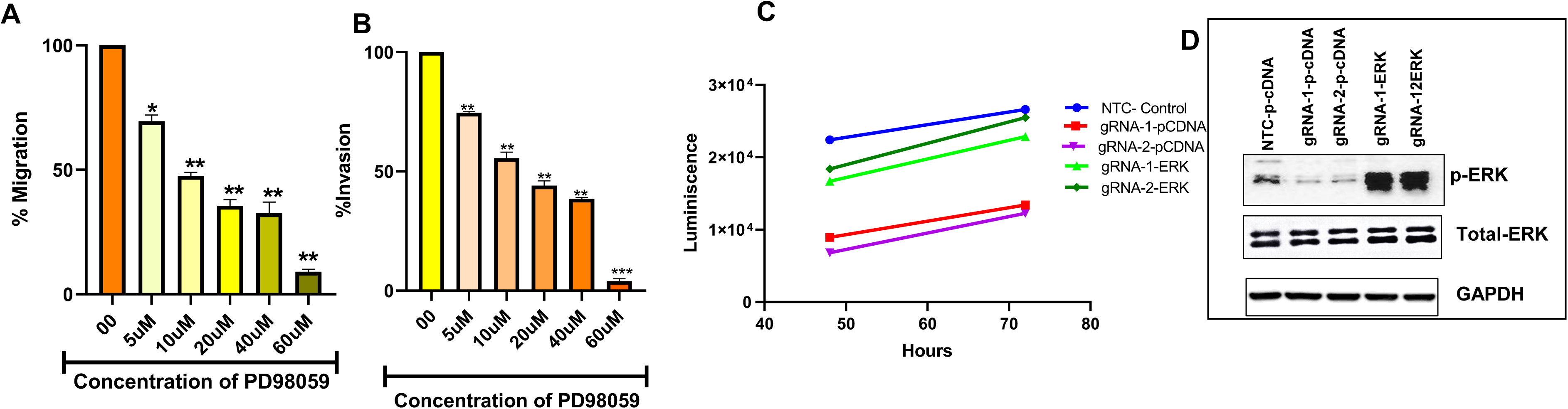
ERK Signaling Modulates EIF3H-Mediated Cell Migration, Invasion, and Proliferation. **(A)**EIF3H-overexpressing (OE) cells treated with increasing concentrations of the ERK inhibitor PD98059 (5-60 µM), followed by migration assay **(B)** EIF3H-OE cells treated with PD98059 (5–60 µM)) and an invasion assay was subsequently performed.**(C)** EIF3H knockdown cells (gRNA-1 and gRNA-2) were transfected with an ERK overexpression plasmid, and cell proliferation was measured at 48 and 72 hours. **(D)** Western blot analysis showing p-ERK levels following ERK plasmid transfection in EIF3H-knockdown cells. GAPDH and Total ERK were used as loading controls.

To validate this interplay, we next overexpressed ERK in EIF3H KD cells and evaluated their proliferative capacity. Remarkably, ERK overexpression rescued the reduced proliferation phenotype of EIF3H-deficient cells (Fig. 5C, D), confirming that ERK acts downstream of EIF3H. Collectively, these findings establish the EIF3H–ERK signaling axis as a critical driver of malignant traits, including proliferation, migration, and invasion. Targeting this axis may offer a promising therapeutic strategy to suppress metastatic progression in breast cancer.

### 5. EIF3H Knockdown Cells Exhibit Enhanced Drug Sensitivity

To assess the role of EIF3H in modulating chemosensitivity, we measured IC₅₀ values and generated dose–response curves for doxorubicin and paclitaxel in EIF3H knockdown (gRNA1 and gRNA2) and NTC cells. For Doxorubicin, the IC₅₀ for NTC was 29 nM (Fig 6A). EIF3H knockdown significantly enhanced drug sensitivity, as indicated by markedly lower IC₅₀ values: 1.67 nM for gRNA1 and 1.433 nM for gRNA2. Similarly, for Paclitaxel, the IC₅₀ for the NTC group was 1.674 nM, while EIF3H knockdown cells exhibited lower values: 0.055 nM (gRNA1) and 0.119 nM (gRNA2) (Fig. 6B). Together, these results demonstrate that EIF3H silencing significantly increases breast cancer cell sensitivity to both doxorubicin and paclitaxel, highlighting the therapeutic potential of targeting EIF3H with small-molecule inhibitors to improve treatment efficacy.

**Figure 6.**
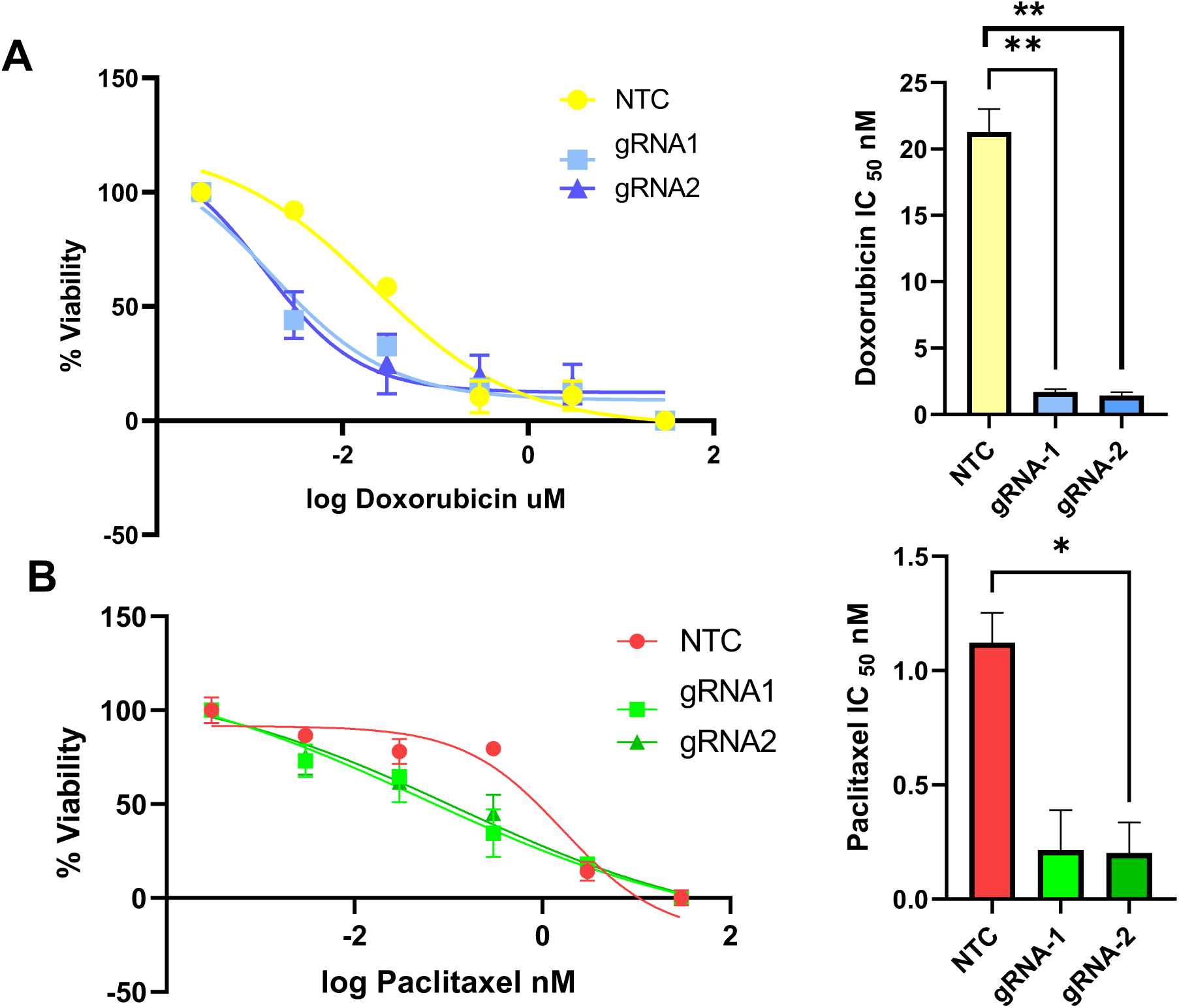
EIF3H Knockdown Sensitizes Cells to Doxorubicin and Paclitaxel: Dose–response curves were generated to determine the IC₅₀ values of **(A)** Doxorubicin and **(B)** Paclitaxel in control (NTC) and EIF3H-knockdown (gRNA-1 and gRNA-2) cells.

## Discussion

Our study identifies EIF3H as a critical regulator of oncogenic signaling and therapeutic response in breast cancer. Using both gain- and loss-of-function approaches in T47D cells, we demonstrate that EIF3H promotes aggressive tumor phenotypes by enhancing cell proliferation, migration, invasion, and chemoresistance. These results corroborate earlier reports of EIF3H overexpression in several cancers, and firmly establish its oncogenic role in breast cancer [10–12]. At the mechanistic level, we identify the ERK pathway as the primary downstream target of EIF3H, wherein EIF3H promotes pERK deubiquitination and stabilization, thereby amplifying MAPK signaling and driving malignant phenotypes (Figure 7).

**Figure 7:**
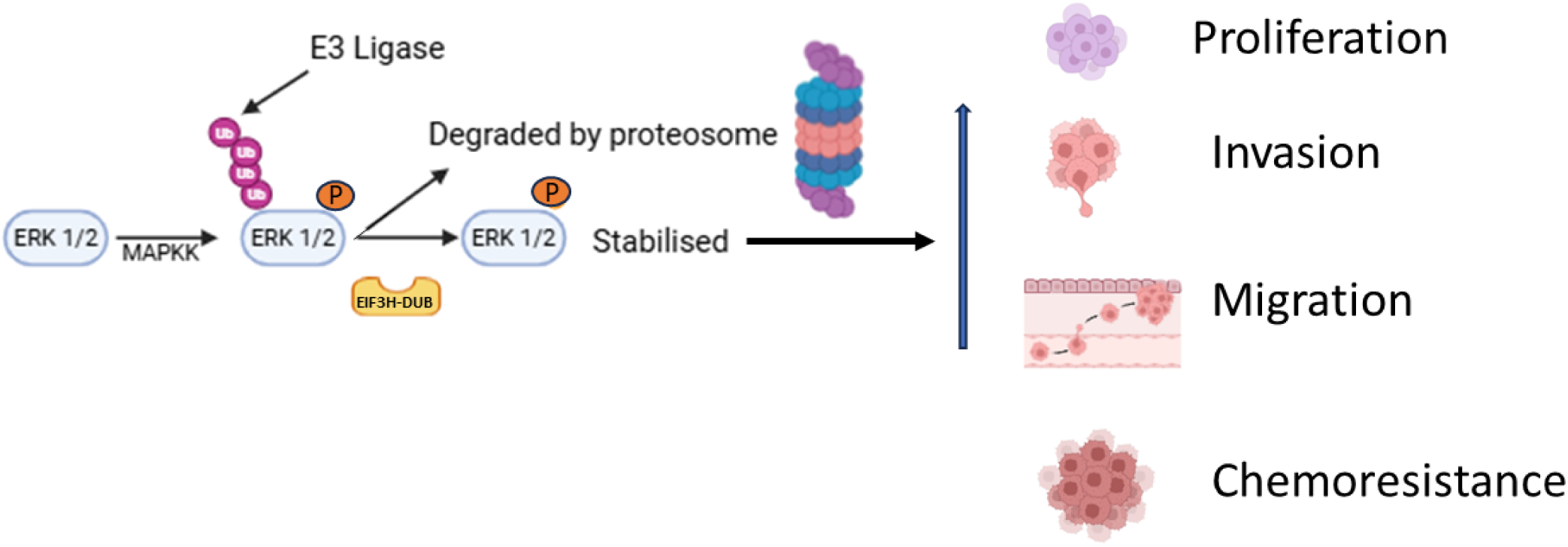
Proposed working model for regulation of pERK by EIF3H in orchestrating breast cancer metastasis.

While ERK regulation through phosphorylation is well established, its regulation via ubiquitination remains comparatively less explored. For instance, MEKK1 not only phosphorylates ERK1/2 but also functions as an E3 ubiquitin ligase that promotes ERK ubiquitination and degradation through its PHD domain [17]. Similarly, ERK1/2 undergoes K63-linked polyubiquitination mediated by TRIM15, with CYLD acting as a counter-regulatory deubiquitinase [20]. In colorectal cancer, EIF3H has been shown to stabilize HAX1 via deubiquitination, thereby activating RAF–MEK–ERK signaling [11]. Building on this, our work is the first to demonstrate that pERK is a direct substrate of EIF3H, which deubiquitinates and stabilizes pERK to sustain ERK-driven tumorigenic processes in breast cancer. This study reveals a previously unrecognized layer of MAPK/ERK pathway regulation, advancing our understanding and highlighting the therapeutic potential of targeting EIF3H to disrupt aberrant ERK signaling in breast cancer.

We further demonstrate that EIF3H knockdown markedly enhances the sensitivity of breast cancer cells to two frontline chemotherapeutic agents, Doxorubicin and Paclitaxel. This effect was evident from the pronounced reduction in IC₅₀ values, indicating that elevated EIF3H expression contributes to chemoresistance. These findings highlight EIF3H as a promising druggable target for overcoming therapeutic resistance. Pharmacological inhibition of EIF3H could potentiate the efficacy of standard chemotherapies, enabling the use of lower drug doses and thereby reducing treatment-associated toxicities. Given the central role of the ERK/MAPK pathway in promoting breast cancer metastasis [16], targeting EIF3H using small molecule inhibitors offers a promising therapeutic strategy to disrupt aberrant ERK signaling and enhance clinical outcomes.

Collectively, our findings identify EIF3H as a multifunctional oncogenic driver in breast cancer. By stabilizing ERK and sustaining MAPK signaling, EIF3H promotes enhanced proliferation, migration, invasion, and resistance to chemotherapy. Given the pivotal role of ERK signaling across multiple malignancies, it is likely that EIF3H exerts similar functions in other ERK-driven cancers, warranting further investigation. Future work should expand the EIF3H interactome to uncover additional substrates within the MAPK cascade or intersecting oncogenic pathways. Moreover, integrating clinical datasets to correlate EIF3H expression with patient outcomes, chemoresistance, and recurrence will be critical for establishing its prognostic value and therapeutic potential.

In conclusion, this study reveals a previously unrecognized mechanistic and therapeutic role of EIF3H in regulating MAPK signaling and driving breast cancer progression. Targeting the EIF3H–ERK axis represents a promising strategy to suppress tumor growth, overcome chemoresistance, and advance the development of more effective, personalized therapies for breast cancer.

## Acknowledgements

We thank Department of Biotechnology, Government of India and Institutional (M.S. Ramaiah University of Applied Sciences) Seed Grant for the financial support.

